# Ants oscillate and scan more in navigation when the visual scene changes

**DOI:** 10.1101/2025.01.13.632872

**Authors:** Sudhakar Deeti, Ken Cheng

## Abstract

Solitarily foraging ants learn to navigate between important locations by comparing their current view with memorized scenes along a familiar route. As desert ants, in particular, travel between their nest and a food source, they establish stable and visually guided routes that guide them without relying on trail pheromones. We investigated how changes in familiar visual scenes affect the navigation of the red honey ant (*Melophorus bagoti*). In Experiment 1, ants were trained to follow a one-way diamond-shaped path to forage and return home. We manipulated scene familiarity by adding a board on their homebound route just before the nest. In Experiment 2, ants were trained to travel a straight path from their nest to a feeder, and we removed the prominent landmarks on the route after they had established a stable route. We predicted that these scene changes would cause the ants to deviate from their usual straight paths, slow down, scan more, and increase their lateral oscillations to gather additional information. Our findings showed that when the familiar scene was changed, ants oscillated more, slowed their speed, and increased scanning bouts, indicating a shift from exploiting known information to more actively exploring and learning new visual cues. These results suggest that scene familiarity plays a crucial role in ant navigation, and changes in their visual environment lead to distinct behavioral adaptations aimed at learning about the new cues.

---

Ants are known to navigate with a toolkit of path integration, landmark-based guidance, and systematic search (Wehner 2020). In path integration, the insect keeps track of the straight-line distance and direction from a starting point, typically its home (Wehner and Srinivasan 2003; Wehner and Wehner 1986). The vector computed by path integration is now believed to be represented within the central complex of the insect brain, specifically in a neural structure that, while not anatomically arranged in a ring, operates as a ring attractor network (Heinze et al. 2018; Lyu et al. 2022). The bump of activity on each part of the neural structure indicates a direction and its amplitude codes distance. In view-based navigation, the ant compares its current view with remembered views at the goal and along the route to the goal (Cheng 2012; Le Moel and Wystrach 2020; Wehner 2003; Wehner and Räber 1979). In using views, ants are now thought to learn scenes to head toward (attraction) and scenes to turn away from (repulsion, Le Moel and Wystrach 2020; Murray et al. 2020). In systematic search, which is frequently needed near the nest when the former two strategies do not take the ant exactly to the nest entrance, the navigator travels in loops that increase in size as the search goes on (Schultheiss et al. 2015; Wehner and Srinivasan 1981).

Loops also feature prominently in learning views. In a number of species that have been studied, a would-be forager first takes loops around its nest before heading off to forage in what are called *learning walks*, three to seven such walks in bull ants (*Myrmecia croslandi*, Jayatilaka et al. 2018) and desert ants (*Cataglyphis bicolor*: Wehner et al. 2004; *C. noda* and *C. fortis*: Fleischmann et al. 2016; Fleischmann et al. 2018; *Melophorus bagoti*: Deeti et al. 2020; Deeti and Cheng 2021a; Deeti et al. 2024a; reviews: Freas et al. 2019; Zeil and Fleischmann 2019). During learning walks, the view learner often stops and looks in various directions, turning on the spot in what are called scanning bouts (Deeti et al. 2023a) or pirouettes (Fleischmann et al. 2017), presumably to learn what the scene looks like in various directions, including the direction to the nest. Ants also scan in bouts on some trips home, especially when experimenters have changed the scene through some manipulation (Deeti et al. 2023a; Wystrach et al. 2014). Presumably, such scans help a navigator to find the best direction to home in.

Even on a familiar route, with or without scanning at the start of the trip, ants do not strike a straight path home. Rather, they oscillate laterally, if only slightly, with the path ‘meandering’ left and right as the ant travels (Clement et al. 2023). Ants, bull ants *M. croslandi* and meat ants *Iridomyrmex purpureus* in the study, oscillate more when the visual scene is unfamiliar than when the scene is familiar.

Oscillatory behaviors are crucial for navigation and widespread across life, from bacteria to animals (Cheng 2022, 2023, 2024). The system or mechanism that produces oscillatory behavior is called an *oscillator*, said to be a basic unit of action (Gallistel 1980), but perhaps is a basic unit of life (Cheng 2022, 2023). Oscillations of effectors that drive movement are common to mobile organisms, but lateral oscillations add a new dimension: the traveler can compare conditions on the two sides as it winds left and right and gather more information about the world as it moves. The lateral oscillatory behaviors of ants in navigation have been described in the past decade (Clement et al. 2023; Collett et al. 2014; Dauzeres-Perez and Wystrach 2024; Lent et al. 2013; Murray et al. 2020). What we want to add to this topic are measures of the characteristics associated with oscillations, frequency, amplitude, and in one case, phase relations between two different oscillations.

We aim to build upon the extant work on lateral oscillations in ants by focusing on one of the extensively studied species of desert ants, the Australian red honey ant *Melophorus bagoti*. We measured the number of bouts of scanning and two kinds of oscillations evident in the videos of moving ants: lateral path oscillations and oscillations or swings of the head from side to side. We defined and measured the frequency and amplitude of these oscillations. For path oscillations, we measured frequencies in time and space. We also examined the (temporal) phase relationship between these two kinds of oscillations, that is, where the head-swing cycles fall on the lateral path oscillations. In addition, we calculated three different measures of straightness to characterize the overall extent of oscillations.

We adopted the hypothesis gathered from the study on meat ants and bull ants (Clement et al. 2023) that in visually unfamiliar circumstances, oscillations would increase. In Experimental 1, we forced ants to travel one-way around a diamond to forage for food and return home. We videotaped the ants from above on their last leg before reaching home. To manipulate scene familiarity, we added a piece of board placed on the side of the homebound track right before the ants reached home. In Experiment 2, we placed prominent boards enroute a straight path from the ants’ nest to a feeder and removed the landmark set up after ants were trained to travel the outbound route. We predicted that the paths with a scene change would be less straight on all straightness measures, that the ants would slow down, that they would scan more, and oscillate more. Our suite of measures of oscillations will let us know in which ways the ants oscillate more. Slowing down, scanning more, and oscillating their heads and their paths more would all serve to gather more information—rather than simply exploiting known information to get home as quickly as possible—a theme that we discuss later.

## MATERIALS AND METHODS

The desert ant *Melophorus bagoti* is common in arid and semi-arid regions in central Australia. The current field-based study was conducted ∼10 km south of Alice Springs, Northern Territory, Australia (23°45.448′ S, 133°52.908′ E). The habitat’s vegetation is dominated by a mosaic of tussocks of the invasive buffel grass (*Pennisetum cenchroides*) interspersed with bushes, *Acacia* spp., and occasional large eucalyptus trees (Deeti and Cheng 2021b; Deeti et al. 2023b; Deeti et al. 2024b). *M. bagoti* form underground monodomous colonies with typically only one entrance. This species is adapted to hot and dry environments, forages during hot periods of summer days, mainly scavenging dead arthropods for a source of protein and gathering sugary plant exudates and seeds as a source of carbohydrates (Muser et al. 2005; Schultheiss and Nooten 2013). There are no local, Territorial or federal ethical requirements to conduct research on ants, and all experiments were non-invasive, with individuals returning to the colony after testing.

### Feeder and channel set up

The experiments took place at two nests of *Melophorus bagoti*. Prior to conducting any experimentation, the grass in the nest area was cleared without altering the rest of the surrounding scenery. We buried a plastic feeder, measuring 15×15×10 cm and featuring a slippery surface, at ground level, 7 m from the nest. The feeder was filled with cookie pieces (Arnott brand) to attract foraging ants.

At Nest 1, the setup consisted of two L-shaped routes, bordered on the sides and forming a diamond, an outbound Nest-to-Feeder Route and an inbound Feeder-to-Nest Route. Both routes shared the same dimensions, with a first leg 5 m in length and 1 m in width, a 90° turn, and a second leg also 5 m in length and 1 m in width (Figure 1). Each route was built using 90cm-long PVC floor tiles with a height of 15 cm. These tiles were buried 5 cm deep in the ground for stability. The PVC floor tiles created a slippery surface for ants, preventing them from climbing over and keeping them on the designated foraging path. The outbound Nest-to-Feeder Route connected the nest to the feeder and the inbound Feeder-to-Nest Route formed the path back home. A one-way foraging circuit was created with three ramps: one at the end of the Nest-to-Feeder Route enabling ants to enter the feeder area (30 cm from the feeder), one at the beginning of the Feeder-to-Nest Route allowing ants to exit the feeder area onto the route home, and the final ramp at the end of the Feeder-to-Nest Route (30 cm from the nest entrance) permitting ants to enter the nest area but preventing them from conducting outbound journeys along this route. The ramps thus served to direct the ants around a one-way circuit counterclockwise around a diamond.

**Figure 1.**
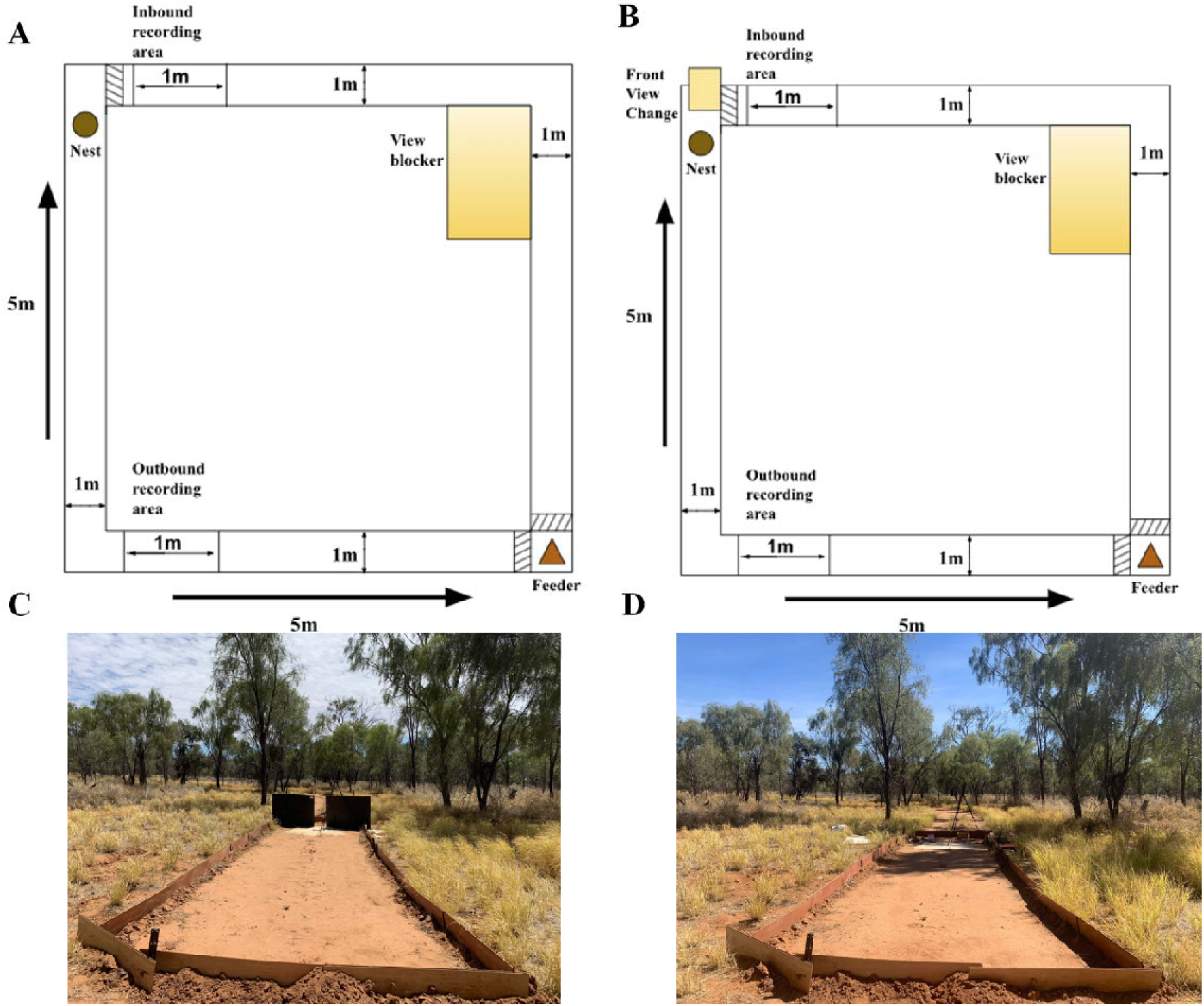
Diagrams showing the diamond setup in Experiment 1, including the recording areas and view blocker (A and B). In B, a plywood board (50 cm × 50 cm) was added just to the right of the recording area. Photos of the track setup in Experiment 2, showing the training condition (C) and the test condition with the wooden boards removed (D)

At Nest 2, foragers were trained to follow a straight outbound route to a feeder located approximately 7 m from the nest and return on the same path. The feeder contained a mix of Arnott-brand cookie pieces and mealworms to attract foraging ants. To guide the ants toward the feeder, a plywood barrier built of the kind of PVC floor tiles that were used for nest 1 was constructed around the route. This barrier spanned 10 m in length and 2 m in width. At a distance of 6 m from the nest, two plywood boards, each measuring 180 × 120 cm, were positioned on the ants’ foraging route with a 50-cm gap between them (Figure 1). This gap intersected the line connecting the nest, the center of the recording area, and the feeder.

To ensure clear visibility of the red ants against the sandy red soil background at both nests, fine white sand was spread on the ground within the recording area. This helped to create a contrast between the ants and the background, making them easily distinguishable in the recorded footage. Once they habituated to the white sand, we observed no notable difference in the behavior of the ants on white sand vs. red soil.

The trajectories of ants collecting food and returning to their nest were recorded at 1 m before the end of the Feeder-to-Nest Route at Nest 1 and 1 m before the feeder, on the outbound route, at Nest 2. To capture the ant paths, a tripod was set up at a height of 1.2 m from the ground. A Sony Handy camera (FDR-AX700) was mounted on the tripod, and the recording was done at a frame rate of 25 frames per second (fps). The recording area had a size of 1 m by 1 m, and the camera had a resolution of 3860 by 2160 pixels. The tripod was left in place throughout experimentation, so that it was consistently a part of the scenery.

### Design

Two experiments were carried out, one at each nest. The two experiments had different set ups and different manipulations, but conceptually, the same kind of comparisons were made. For each ant, behaviors captured on two different video recordings were compared. Recordings were made on the trip just before a scene change and the first trip with the scene change. This allowed before–after comparisons to be made with regard to each scene change.

### Video analysis

To track and analyze behavior, we used a custom trained DeepLabCut™ pose-estimation model to annotate the positions of the ants in each frame of video. We tracked 2 body parts, the front of the head and the middle of the thorax. We labelled 500 frames for training, We used a resnet50 pretrained backbone (trained on imagenet). We trained for 50,000 iterations and selected snapshot 3 based on the average pixel error on the test dataset. To ensure that this trained model was effective, we viewed some videos annotated by the trained model and also viewed the plotted predictions for every treatment replicate. We used a confidence cutoff of 0.6. All coordinates are plotted as x–y coordinates of the video frame. We used the position of the ant in the first detected frame as the 0,0 location. At Nest 1, y coordinates point in the direction of the nest from the feeder, whereas at the Nest 2, they indicate the feeder-to-nest direction.

To smooth out the data and reduce noise, we first identified each stationary frame (stop threshold < 0.4 mm/s) and then ignored and excluded these points from the data. Sequences of points less than this distance apart are coalesced into a single point to produce a path that ‘ignored’ stops. We then computed a 5-point moving average of the x and y coordinates in head and thorax positions in each trajectory to reduce some inevitable noise stemming from the camera and coding system. Other than this smoothing, we did not transform, recode, or remodel the data any further, basing all measures on the smoothed x–y coordinates. The measures of oscillations were calculated from the x–y coordinates and not from a neural-net or any other kind of model applied to the coordinates. Paths were characterized and visualized in R (version 4.2.1; R Core Team, 2020) using the packages trajr (McLean and Skowron Volponi 2018) and Durga (Khan and McLean 2023; Deeti et al., 2024c; Deeti et al., 2024d).

### Summary statistics

We used the extracted frame-by-frame coordinates to calculate the final heading direction, speed, orientation direction, orientation angular velocity and path characteristics. The final heading direction was the vector from the thorax to the head on the final frame of a video record. This vector represented the orientation of each ant at that specific moment. To understand how quickly and directly the ants moved at the recording locations, we calculated the speed of the workers along with several characteristics of the entire trajectory. These path characteristics were based on the positions of the thorax across frames. For each ant, speed was measured by the magnitude of an ant’s velocity, calculated as path length covered by the entire trajectory divided by the duration of the trip, excluding the stopping durations. We measured the speed at which ants changed their orientation direction by observing how quickly the thorax–head direction changed over time. As ants walk, they turn their heads left and right in oscillation.

For the entire path, we calculated three indices of straightness: *path straightness, sinuosity*, and 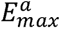, each of which relates to the directness of navigation toward a destination. *Straightness* of the trajectory is computed as the ratio of the straight-line distance (*D*) traversed by the recorded trajectory to the overall length (*L*) of the path (Batschelet, 1981; Deeti et al. 2023c; Islam et al. 2021; Islam et al. 2023). The formula for straightness, *D/L*, ranges from 0 to 1 (as *D* ≤ *L*), with larger values indicating straighter paths, while smaller values indicate more curved or convoluted paths. *Sinuosity* is an estimate of the tortuosity or extent of moment-to-moment winding of the recorded path, calculated as 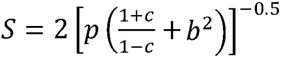, where *p* is the mean step length, *c* is the mean cosine of turning angles and *b* is the coefficient of variation of the step length. A trajectory *step* refers to the movement of the animal’s thorax position recorded in consecutive video frames. Step lengths are the Euclidean distances between consecutive thorax positions along a path, while turning angle refers to the change in orientation direction between two consecutive steps. Sinuosity varies between 0 (straight path) and 1 (extremely curved path) (Benhamou 2004; Lionetti et al. 2023). The maximum expected displacement of a path 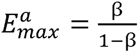, where β is the mean cosine of turning angles, is a dimensionless value expressed as a function of the number of steps. It aligns with the intuitive meaning of travelling straight (Cheung et al. 2007). Larger maximum-expected-displacement values indicate straighter paths with a greater displacement in the trajectory, while smaller values suggest more localized or constrained movement. Ants frequently displayed a series of successive fixations in different directions by rotating on the spot at one location, known as a “scanning bout” (Deeti et al. 2023a; Islam et al. 2022). For each ant in each condition, we extracted the number of scanning bouts and the scanning-bout durations, from the start of a scanning bout until the ant started walking again.

To characterize oscillations of paths, we calculated amplitude, temporal frequency, and spatial frequency. Initially, we segmented ‘wiggles’ by identifying inflection points on the path. Points of thorax positions turning in one direction formed a wiggle. We specified a minimum of three turns in the same direction for a wiggle, that is, one or two turns in the opposite direction did not count as a change of direction. Each point just before a change of direction was an inflection point. The amplitude of each wiggle was the point with the maximum distance from the straight line connecting the end points of the wiggle, tallied in centimeters. The spatial period of the wiggle was calculated from the length of the straight line between the inflection points defining a wiggle and doubling it (one wiggle being half a cycle), also in centimeters. The spatial frequency was then derived by taking the inverse of the spatial period (cycles/cm). To calculate the temporal period of a wiggle, we counted frames between inflection points and doubled the count. The number of frame changes was translated into seconds by dividing by the frame rate of 25 f/s. The temporal frequency was the inverse of the temporal period (cycles/s). We excluded the first and last wiggles due to unknown inflection points beyond the data range. For each of the remaining wiggles, we calculated for each ant the average and standard deviation in frequencies and amplitudes across videotaped segments of that ant, plotting distributions of means and standard deviations across ants for visualization.

For orientations, each change in direction of heading from left to right or from right to left was taken as an inflection point, again with a minimum sequence length of 3 frames. Temporal frequencies and amplitudes, the latter measured as degrees of orientation change, could then be calculated in an analogous fashion to the calculations for path-oscillatory characteristics. Spatial frequencies do not make any sense in the context of head orientations, as position in space does not form a defining role in the thorax–head direction.

With two separate oscillations, a third parameter can be calculated, that of the phase relationship between head orientation and path oscillations. Head orientation changed faster than the path ‘wiggle’, so that we plotted the former on the latter cycle. Double wiggles of the path were represented as a circle with 0 representing the start of the turn to the left. Angles thus represented proportions of a period. We plotted the starts of head-orientation cycles (start of turn to the left) on this circle, with the angles increasing clockwise, combining all ants and all segments. We repeated this exercise with the start of a right turn on both cycles as 0. Because of unevenness in oscillations, if the start of the left turn is 0°, the start of the subsequent right turn is not exactly at the midway point in time or 180° on a circular graph but ranged from −27° to 140°.

### Experiment 1

#### Nest-1 Control

To investigate the impact of changes in the learned view panorama on the navigational behavior of *Melophorus bagoti*, we first conducted a training phase to familiarize the ants with the views encountered while traveling through the channels. The foraging ants (N =15) were allowed to learn the route from the nest to the feeder and from the feeder back to the nest on their own. They usually learned to navigate from the nest to the feeder and back to the nest within a day. Once an ant successfully located the feeder and returned close to the nest, we captured it and applied a specific color code to its body (Citadel brand). This painting allowed us to identify individual ants. The painted ants were then observed, and they were given the opportunity to learn and familiarize themselves with the route for 3 days. If any ant failed to complete at least 5 trials during the three-day period, we excluded them from the experiment. At the end of the third day (14:00 to 17:00), we recorded one trip of each trained ant at the recording location for this Control1 condition. In a few instances when two ants appeared simultaneously, the second ant was temporarily intercepted before entering the final segment of the Feeder-to-Nest channel. These intercepted ants were not included in the study.

#### + Board

In the +Board condition (N =15), the ants that had completed the control condition were subjected to a change in their visual environment on the fourth day. The foragers started the activity at ∼9:00 in the morning and activity reached a maximum from ∼10:00. Once, they found the food at the feeder, they continued foraging actively throughout the day. At the start of day 4, we implemented the view change at Nest 1: a brown plywood board measuring 50 cm in width and 50 cm in height was placed after the exit ramp at the end of the Feeder-to-Nest Route (Figure 1). This plywood board altered the ants’ visual scene by blocking a part of their previously familiar view. The behavior of the ants encountering the view change was observed and recorded with the same video set up as previously discussed. Each ant was recorded only once at the designated location in this condition. Once recorded, ants were captured within 5 cm of the nest entrance and marked with a black color on their abdomen. This marking prevented the same ant from being recorded multiple times.

#### Control comparisons

To test if the first trip of the day produced different behaviors in the ants even without any view changes, we compared the last recorded trip of each subject ant on day 2 with its first trip on day 3. These trials followed the procedures of the Control1 and +Board conditions except that no board was added on day 3.

### Experiment 2

#### Nest-2 Control

In this condition (N = 15), foragers were given the opportunity to learn the route to the feeder 7 m from the nest on a straight route. Upon their initial arrival at the feeder on the first day, foragers were marked with a distinctive color code (Citadel brand) on the abdomen and then released at the feeder. The painted ants were then given three days to learn the route and the landmarks along the way to feeder. As with Experiment 1, at the end of the third day (14:00 to 17:00), we recorded an outbound trip of each trained ant at the recording location for this Control2 condition. Any ant that failed to complete at least 5 trials during the three-day period was excluded from the experiment.

#### – Board

In this board-removal manipulation (N =15), the ants that had completed the control condition were tested with a change in their visual environment on the fourth day. Before the ants’ first trip of the day, the two boards that had been on the route to the feeder were removed. The behavior of the ants encountering the route, now without boards, was observed and recorded with the same video set up as before. Each ant was recorded only once. Once recorded, an ant was captured at the feeder and marked with a black color on its abdomen to prevent multiple testing of the same ant.

#### Control comparisons

To test if the first trip of the day produced different behaviors in the ants even without any view changes, we again compared the last recorded trip of each subject ant on day 2 with its first trip on day 3. These trials followed the procedures of the Control2 and –Board conditions except that the boards were still present on day 3.

### Statistical analysis

The experiments conducted at the two nests were analyzed using paired Welch’s t-tests, which are suitable for groups with unequal variances. In each experiment, the control and test data were paired, and the magnitude of change in each statistical unit was estimated. We employed models to compare the effect of view change on the following variables: speed, orientation change velocity, path straightness, sinuosity, 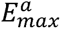, number of scanning bouts, duration of bouts, orientation amplitude mean, orientation amplitude standard deviation (SD within each individual across cycles in the video record), orientation temporal frequency mean, within-individual orientation temporal frequency SD, path amplitude mean, within-individual path amplitude SD, path temporal frequency mean, within-individual path temporal frequency SD, path spatial frequency mean, and within-individual path spatial frequency SD, with alpha set at P=0.01. We predicted that speed, path straightness, and 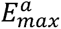 would decrease, while sinuosity, the number of scanning bouts, and scanning bout durations would increase in the test conditions. For oscillatory behaviors, we predicted that frequencies or amplitudes would increase, but we have no specific predictions. We had no predictions regarding orientation change velocity.

### Data Transparency and Openness

Our data and R scripts for both Experiment 1 and Experiment 2 are made publicly available at (https://osf.io/6wk7a/files/osfstorage). The predictions and methodologies for both experiments in this study were not pre-registered.

## RESULTS

To investigate the potential impact of a view change on the learned routes of foragers, we recorded and visualized the path trajectories of ants while returning from a feeder to their nest as well as going to the feeder from the nest. We captured data in four scenarios: nest-1 ants before the view change while returning from the feeder (Control1) and after the view change during the return journey from the feeder to the nest (+Board) (see Figure 1), nest-2 ants while en route to the feeder (Control2) and after the view change (removal of boards, – Board). As a general observation that was not the focus of this study, ants in all conditions navigated to their goals without exception; the experimental changes did not affect navigational success. Compared to the Control conditions, however, ants in the +Board and – Board conditions exhibited slower speeds, more meander (Figure 2), and an increased number of scans. Specifically, 12 out of 15 ants in the +Board condition and all 15 ants in the –Board condition performed scans, with ants conducting between 1 to 18 scans per meter in test conditions. In contrast, the majority of ants in the Control conditions did not perform any scans. View changes also altered oscillatory characteristics. In our control comparisons (end of day 2 vs. start of day 3), we found no significant differences in both experiments between day 2 and day 3 in any measure of straightness, in speed, and in orientation angular velocity (S. Figure 1). Hence, we will ignore these control comparisons and focus on comparisons with experimental conditions of view changes (S. Table 1).

**Figure 2.**
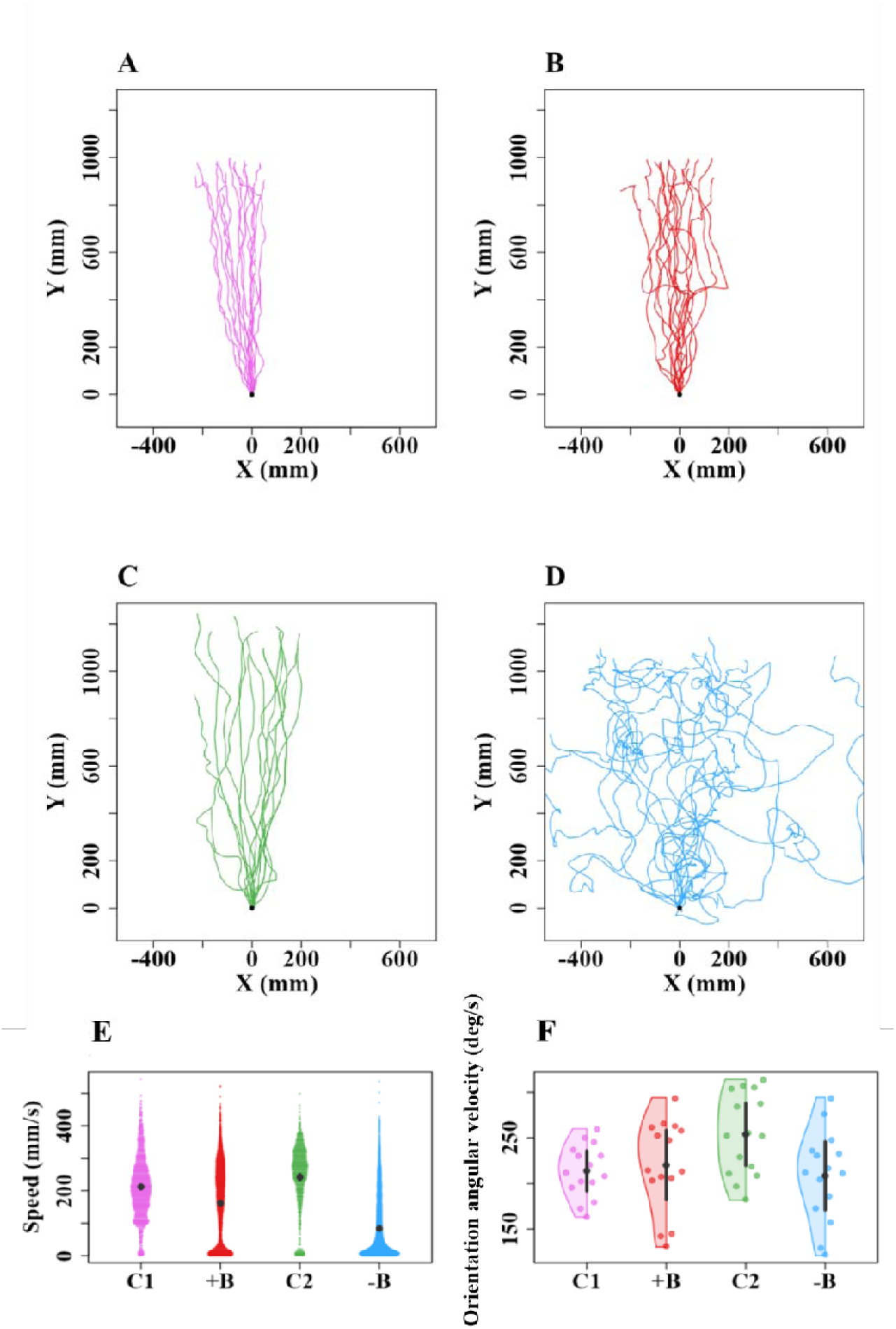
Trajectories, speed, and head orientation of foragers in different conditions. Trajectory plots depicting the paths of ants under different experimental conditions. (A) Control1: Ants in Experiment 1 returning to the nest after foraging with food. (B) +board: Ants exposed to a view change during their homebound run with food. (C) Control2: Ants in Experiment 2 departing from the nest for foraging. (D) –board: Ants exposed to removal board during their foraging runs toward the feeder. Each trajectory plot represents the movement path of an individual ant, plotted over time. The y coordinates represent the spatial position of the ants toward the goal direction. The coordinates (0,0) represent the starting frame of the recording. Comparison of speed and orientation angular velocity of foragers in different conditions (E, F). The violin plots show the mean speed of ants across their entire trajectory (E). The half-violin plots show the distribution of bootstrapped differences of orientation-change velocity of ants (F). The solid dot shows mean, while the vertical bar shows 95% confidence interval of the mean.

When the view changed, with a new landmark on their way home or missing landmarks on their way to the feeder, the ants slowed down, meandered more, looked around more often, and stayed in the recording area longer compared to the normal conditions (Figure 2). In speed, the Welch’s t-tests revealed significant differences in both comparisons (C1 vs. +B: *t* = –5.4, *df* = 14.7, *p* = 0.00007; C2 vs. –B: *t* = –9.2, *df* = 24.6, *p* = 0.00001). With the change in view, the foragers’ orientation angular velocity decreased in magnitude (Figure. 2F). Welch’s t-tests, however, showed no significant differences in orientation angular velocity between Control and view-change conditions (C1 vs. +B: *t* = –0.5, *df* = 27.7, *p* = 0.5; C2 vs. –B: *t* = – 2.6, *df* = 25.9, *p* = 0.04).

In the view-change conditions, the ants’ paths showed increased *sinuosity* and decreased *straightness*, indicating more curved and winding trajectories (Figure 3). Additionally, the *E^a^_max_* values were lower, suggesting greater deviation from their expected navigation path. *Sinuosity* increased in the view-change conditions (Figure 3A). The Welch’s t-tests, however, showed a trend with the board addition and a significant difference with board removal (C1 vs. +B: *t* = –2.4, *df* = 17.7, *p* = 0.02; C2 vs. –B: *t* = –5.7, *df* = 25.8, *p* = 0.00003). 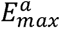 was lower in the view-change conditions, with the ants having a smaller amount of displacement per unit length travelled compared to the control conditions (Figure 3B). The Welch’s t-tests, however, showed a significant difference in Experiment 2 but not in Experiment 1 (C1 vs. +B: *t* = 1.2, *df* = 25.8, *p* = 0.2; C2 vs. –B: *t* = 3.6, *df* = 14, *p* = 0.002). With *straightness*, the view change led to a significantly higher magnitude of path deviation from a straight-line path compared to paths in the Control conditions (C1 vs. +B: *t* = 4.7, *df* = 20.2, *p* = 0.0001; C2 vs. –B: *t* = 16.2, *df* = 17.2, *p* = 0.00006) (Figure 3C).

**Figure 3.**
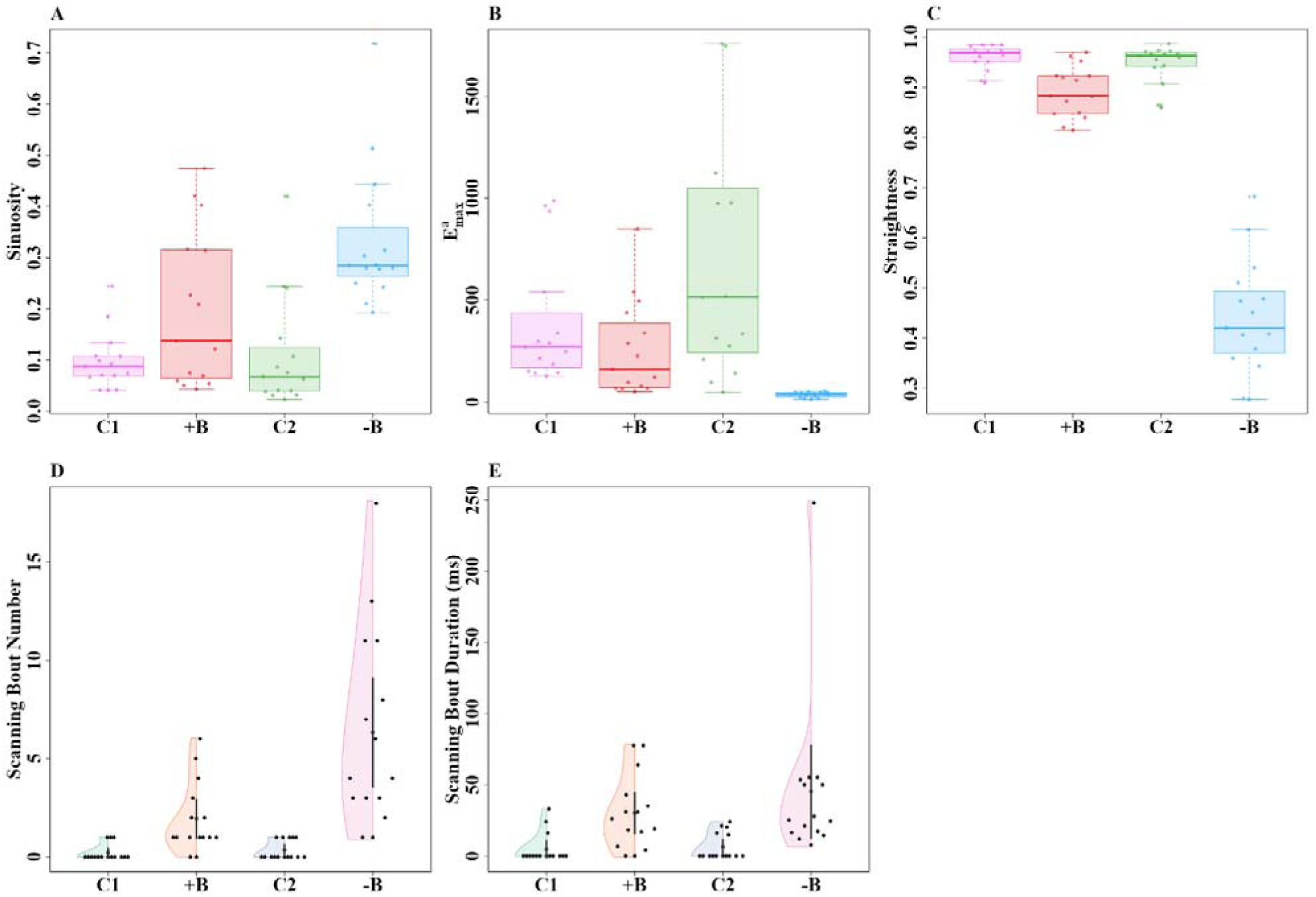
Path characteristics and scanning of ants under different experimental conditions. Shown are the path characteristics of *Sinuosity* (A), *E^a^max* (B) and *Straightness* (C) of trajectories, the number of scanning bouts (D) and scanning-bout duration (E) in the different conditions. Box plots display the median (line inside the box), interquartile range (box), and extreme values excluding outliers (whiskers). Individual data points are shown as dots. On the x axis, C1, +B, C2 and –B denotes the Control1, board addition, Control2, and board removal conditions, respectively. The half-violin plots show the distribution of bootstrapped differences; the solid dot shows the mean, while the vertical bar shows the 95% confidence interval of the mean.

View change increased scanning in foragers. In Control recordings, i.e. those without any scene change, three ants (3/15) in Control1 and four ants (4/15) in Control2 performed a single scan each (Figure 3D), while the rest did not perform any scans at all in the recording area. In contrast, in +B, six ants scanned at least twice and all of the ants at least once in the recording area and in –B, thirteen foragers performed more than two scans and all ants performed at least one scan (Figure 3D). The Welch’s t-tests revealed significant differences in both comparisons, (C1 vs. +B: *t* = –3.6, *df* = 15.4, *p* = 0.002; C2 vs. –B: *t* = –4.6, *df* = 14.2, *p* = 0.0003). The durations of scanning bouts also increased significantly with view changes (Figure 3E; C1 vs. +B: *t* = –3.4, *df* = 18.1, *p* = 0.002; C2 vs. –B: *t* = –2.5, *df* = 14.1, *p* = 0.01).

Ants oscillated their orientation as they walked, swinging their heads left and right. We analyzed the amplitude and temporal frequency of these oscillations in the two view-change and two control conditions. In the view-change conditions, head orientation amplitude mean and SD were higher compared to controls (Figures 4A and 4B). Together with slower forward movement with view change, ants also showed slower head movements, resulting in a lower temporal frequency of orientation oscillations (S. Figure 2A). Welch’s t-tests showed borderline significance in Amplitude mean with board addition (C1 vs. +B: *t* = –2.5, *df* = 16.4, *p* = 0.01) and a significant difference with board removal (C2 vs. –B: *t* = –3.9, *df* = 24.6 *p* = 0.0005). Amplitude SDs did not increase significantly with board addition (C1 vs. +B: *t* = –1.8 *df* = 19.8, *p* = 0.02) but increased significantly with board removal (C2 vs. –B: *t* = –5.2, *df* = 26.3, *p* = 0.00002). The paired t-tests did not find significant differences in Temporal frequency in board-addition condition (C1 vs. +B: *t* = 0.6, *df* = 25.8, *p* = 0.4) but found a significant decrease with board removal (C2 vs. –B: *t* = 4.2, *df* = 25.6, *p* = 0.0002). The SD of temporal frequency (S. Figure 2B) did not differ significantly with board addition (C1 vs. +B: *t* = –0.2, *df* = 27.9, *p* = 0.7) or board removal (C2 vs. –B: *t* = 1.5, *df* = 23.9, *p* = 0.13).

**Figure 4.**
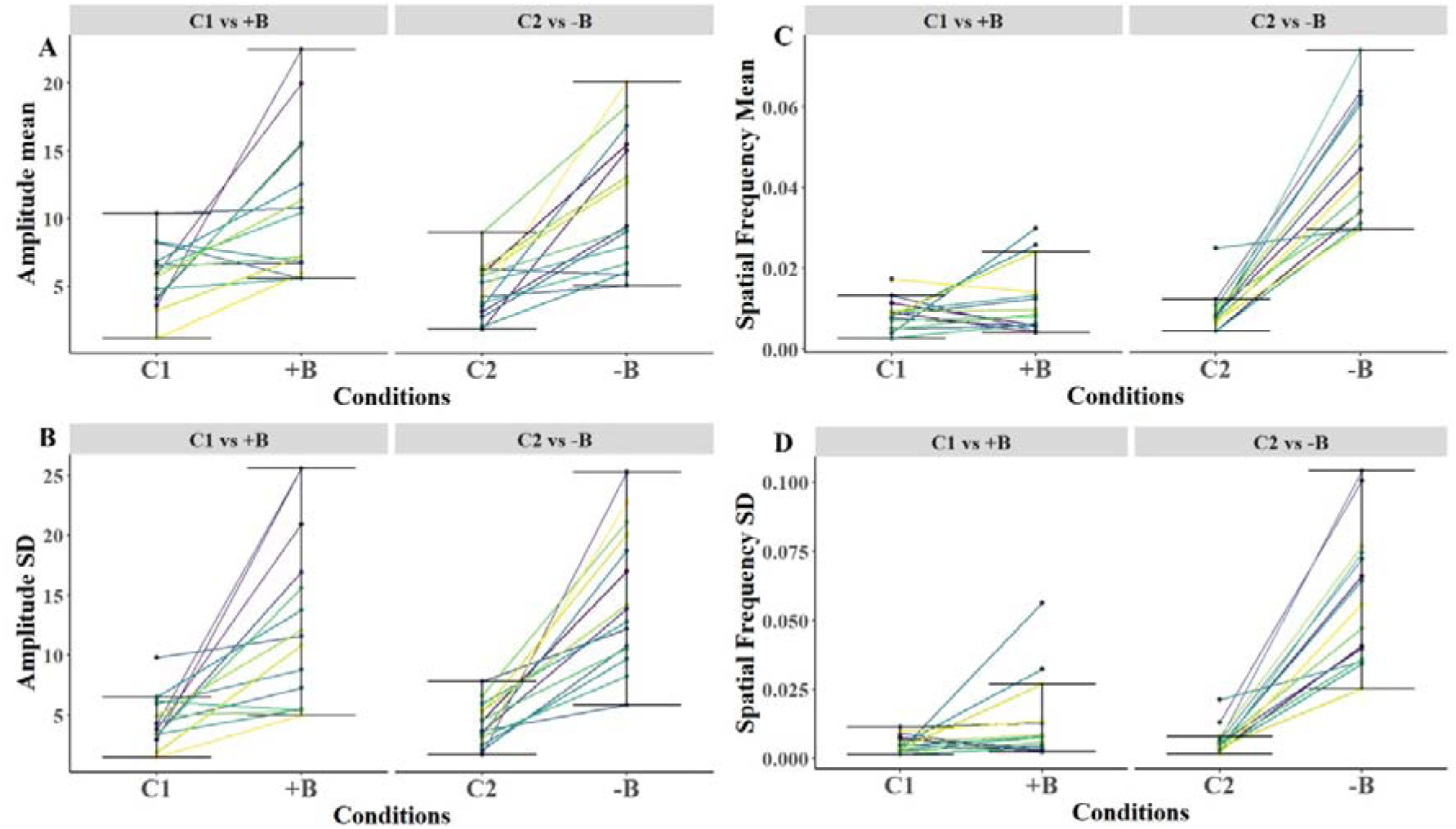
Amplitude of orientation oscillations and Spatial frequency of path oscillations. The plots show the effects of scene change on Amplitude mean (A), Amplitude SD (B) of orientation oscillations, Spatial frequency mean (C) and Spatial frequency SD (D) of path oscillations from the control to the test conditions. The SDs are the SDs across all the ‘wiggles’ of an individual. On the x axis, C1, +B, C2, and –B denote the Control1, board addition, Control2, and board removal conditions, respectively. Units are degrees for amplitude and cycles per cm for spatial frequency.

We then examined the effects of view change on path oscillations, comparing means and SDs in Amplitude, Temporal frequency, and Spatial frequency (Figure 4, S. Figure 2). The paired t-test on Amplitude showed no significant differences in the two comparisons (C1 vs. +B: *t* = –0.4, *df* = 25.9, *p* = 0.6; C2 vs. –B: *t* = –2.1, *df* = 27.2, *p* = 0.04) (S. Figure 2C). Amplitude SD showed inconsistent effects across comparisons in the t-tests (C1 vs. +B: *t* = – 0.9, *df* = 24.3, *p* = 0.36; C2 vs. –B: *t* = –3.9, *df* = 27.1, *p* = 0.0005) (S. Figure 2D). In Spatial frequency mean, the paired t-tests showed inconsistent results in two group comparisons (C1 vs. +B: *t* = –1.3, *df* = 20.6, *p* = 0.19; C2 vs. –B: *t* = –9.5, *df* = 17.2, *p* = 0.00001) (Figure 4C). In Spatial frequency SD as well, the t-tests revealed inconsistent differences in group comparisons (C1 vs. +B: *t* = –0.5, *df* = 18.5, *p* = 0.5; C2 vs. –B: *t* = –8.01, *df* = 15.1, *p* = 0.00007) (Figure 4D). In Temporal frequency means, results did not differ significantly across comparisons (C1 vs. +B: *t* = 0.8, *df* = 25.3, *p* = 0.3; C2 vs. –B: *t* = 1.05, *df* = 25.4, *p* = 0.3) (S. Figure 2E). Temporal frequency SD also did not show significant differences in any comparison (C1 vs. +B: *t* = 0.1, *df* = 27.9, *p* = 0.8; C2 vs. –B: *t* = 0.7, *df* = 18.5, *p* = 0.4) (S. Figure 2F).

In the measures of oscillatory characteristics, means and SD correlate positively across individuals in all the conditions, showing a form of Weber’s law across individuals, in that the SDs of a measurement increase linearly with the mean (Figure 5). Assuming a 50% chance level, the random chance that 20 correlations all turn out positive is 2^−19^, two-tailed. Amplitudes of orientation oscillations and path oscillations, along with spatial frequency in path oscillations, showed significant correlations in all conditions. (Table 1).

**Figure 5.**
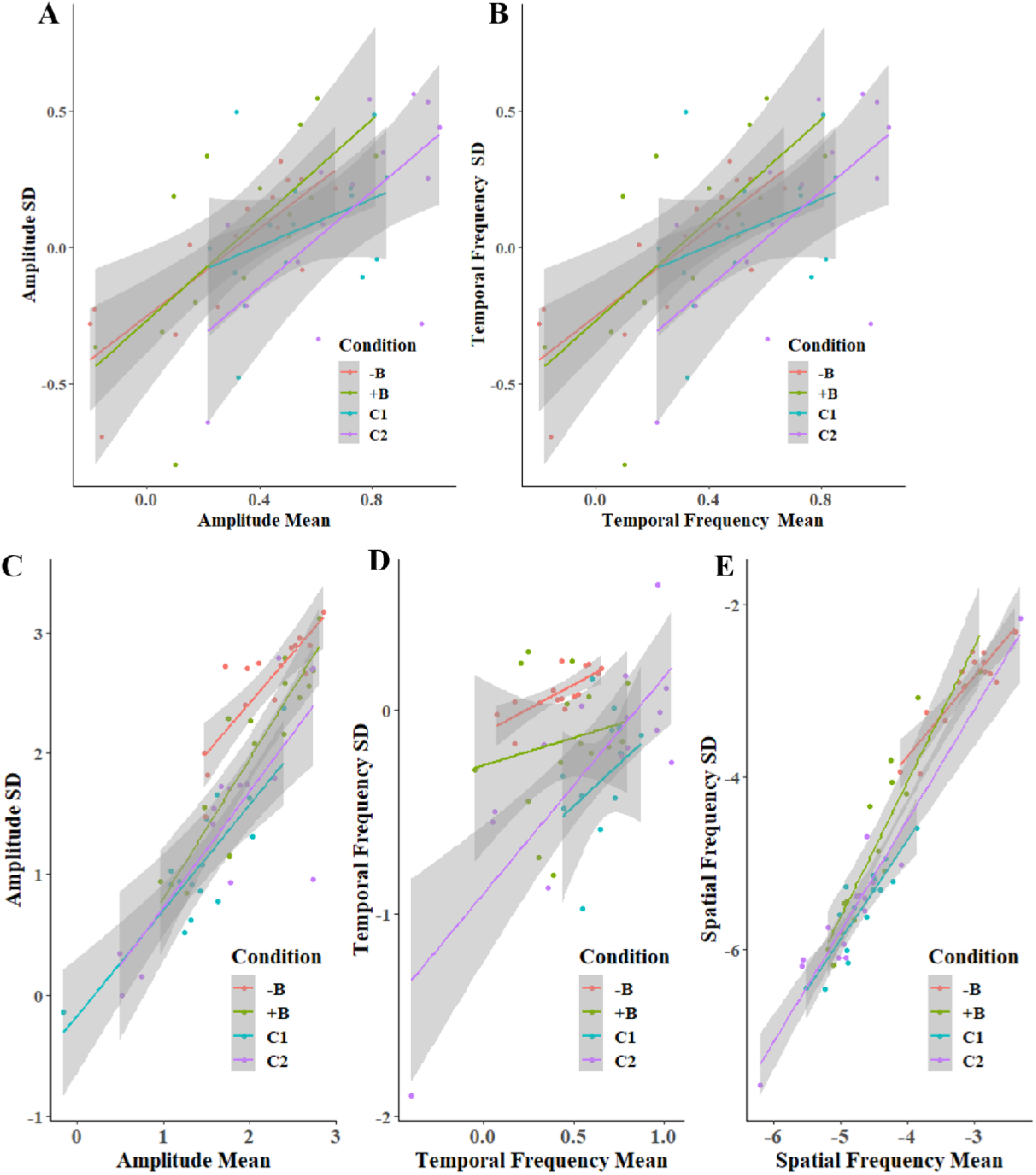
The correlation between means and SDs of oscillation parameters. The scatter plots show the correlations for each condition between the means and within-individual standard deviations, with gray bands marking the 95% confidence intervals. C1, +B and C2 and –B denote the Control1, board addition, Control2, and board removal conditions, respectively. For better visualization values are transformed to log scale on both axes.

To investigate the phase relationship between path oscillations and orientation oscillations, we plotted the initiation points of orientation cycles onto the cycles of path oscillations, designating 0° as the start of left and right turns in two separate analyses (S. Figure. 3A and 3B). The circular distributions revealed that, in general, the initiation points of head turns were scattered and uniformly distributed throughout the path-oscillation cycle in both cases, giving no evidence for phase-coupling between the two types of cycles.

## DISCUSSION

Our experimental view changes did not affect navigational success in the desert ants, as all of them arrived successfully at their goals. But the addition or especially the removal of boards along the route changed the walking characteristics and oscillations of the ants. The ants slowed down, scanned more and for longer durations, and their paths were less straight. The ants engaged in at least three types of oscillations: the legs lifted and planted in cycles, the head swung left and right, and the path wiggled left and right. With view changes, the head swung more to the left and right, increasing amplitudes of orientation oscillations. With board removal, although not with board addition, spatial frequency of path oscillations increased. These results demonstrate the existence of oscillators at work in navigation (Clement et al. 2023), with navigational demands adjusting the working of oscillators (Cheng 2022, 2023). The ants’ reactions to view changes and the workings of oscillators form main themes for this discussion.

### Reactions to view changes

The suite of behavioral changes in navigation in the face of a scene change makes sense as strategies to gather more visual information about the environment. An ant traveling with slower speed than usual, with more meander, more turning of the head, and more stopping to scan for longer would be looking at the environment in more directions and from more places than usual. With a scene change, the ants searched and likely learned the new information. As ants showed oscillations, especially head turns, even on the paths well traveled (in the Control conditions), they are engaging in some information seeking in all navigation. The balance between exploration, the seeking of potentially new information, and exploitation, striding off the well-known route (Clement et al. 2023), tips to more exploration with view changes. But the trade-off between exploitation and exploration, first noted to our knowledge by Kramer and Weary (1991), is still in play in routine runs home or in ‘control’ conditions, testifying to the importance of exploration (Brembs 2011, 2021). The ‘imperfections’ or variability in the oscillatory cycles inject variability in behavior, an important component in exploring.

With regard to other species, view changes on a well-traveled route reduced night-active bull ants’ navigational efficiency significantly (Islam et al. 2020; Islam et al. 2021; Narendra and Ramirez-Esquivel 2017). While the view changes observed in Islam et al.’s (2020, 2021) studies were large, the view change in Narendra and Ramirez-Esquivel’s (2017) study was subtle, consisting in the felling of three trees in a forested surround. The authors used arrows in their illustrative figure to point out the change caused by the tree felling; the difference was not easy for readers to spot. The red honey ants, in contrast, cope with view changes with aplomb, continuing to navigate successfully after being raised a meter off the ground (Julle-Daniere et al. 2014) or with one side of the usual view blocked (Schwarz et al. 2014). None of these studies examined oscillatory characteristics. The bases of these differences in reactions are unclear. Species differences and night vs. day travel both come to mind as possibilities.

Our study found that board removal on the outbound route had larger effects than board addition on the inbound route. Although this is not a focus of the current study, we believe that the difference stems from the much larger view change in the board-removal manipulations. In Experiment 2, the boards on the outbound route were a good deal larger than that in Experiment 1, and two of them stood right on the path to the goal (feeder). Future studies might examine ‘dose-response’ curves in reactions to visual changes.

Previous studies on desert ants have found an increase in scanning or meandering or both when matters are not usual in travel (Freas et al. 2022; Wystrach et al. 2014, 2019, 2020). When views are unusual at the start of the journey home, ants scan more (Wystrach et al. 2014). When an ant had been picked up near the end of its journey home and placed back on the route, the traveler still heads in the home direction, but scans and meanders more (Wystrach et al. 2019; in bull ants, Deeti et al. 2023c; Lionetti et al. 2024). When an ant had encountered a trap that delayed its trip home, it scans more as it nears the location of the trap on the subsequent trips (Freas et al. 2022; Wystrach et al. 2020). We theorize that all these manipulations induce something equivalent to uncertainty in the ants. Whatever proxy measures for uncertainty the ants are relying on, the outcome can be summarized as more exploration under conditions we experimenters would call uncertain, again reflecting the trade-off between exploration and exploitation (Brembs 2011, 2021; Clement et al. 2023).

Finally on this topic, we have replicated the pattern of reactions to view changes in a separate study on nest 2 for a different purpose (Deeti et al., submitted). Changing the color of the boards on the way to the feeder also led to increased scanning and meander, higher spatial frequency of path oscillations, and higher amplitude of head oscillations.

### Oscillators and oscillations

We found path oscillations and head oscillations, the former replicating similar findings in day-active Australian bull ants (*Myrmecia croslandi*) and meat ants (*Iridomyrmex purpureus*; Clement et al. 2023), in wood ants (*Formica rufa*; Collett et al. 2014; Lent et al. 2013), and in desert ants *Cataglyphis velox* (Dauzeres-Perez and Wystrach 2024). Path oscillations may thus be common in ants, and it is also found in many other insects (Clement et al., 2023; Namiki and Kanzaki, 2016). Whereas Clement et al. (2023) used sophisticated Fourier analysis to extract characteristics of oscillations, our approach was to base calculations on the x–y coordinates extracted from the video records. Aside from some smoothing, also used in the Clement et al. (2023) study, our definitions of frequency and amplitude used the spreadsheet data without further recoding or modeling. That such measures can be readily measured based on x–y coordinates of the locations of chosen points on the ants’ body testifies to the robustness of head and path oscillations in ant navigation.

Navigational servomechanisms operate with intrinsic oscillators in animals and even non-neural organisms (Cheng, 2022, 2023), with the oscillators responsible for generating periodic movements of effectors that drive locomotion known as oscillations. Oscillators come with costs. Compared with random-rate processes in behavior, which might rely on noise in the systems generating behavior (Berg and Brown 1972; Deeti et al. 2023a; Deeti et al. 2024e; Scharf et al. 1998), a dedicated mechanism is needed to orchestrate oscillations, presumably relying on neurons in animals. With head and path oscillations seemingly independent in our data set, two such systems are needed. Both neurons and wiring come with hefty costs (Sterling and Laughlin 2015). An additional cost is the time taken to weave left and right, compared with a straight course. These basic units of action (Gallistel 1980)— oscillators generating lateral oscillations—so common in insects (Clement et al. 2023; Namiki and Kanzaki 2016) and other organisms (Cheng 2022, 2023) must provide benefits.

Our discussion of reactions to visual changes has already identified a key benefit: exploration. Frequency and amplitude can be considered as ‘free’ parameters that can be adjusted servomechanistically to serve navigational functions. Variability in behavior in general has other benefits than exploration, including evading predators and searching a ‘problem space’ for optimal solutions to a behavioral challenge (Brembs 2011, 2021).

Oscillations have yet another benefit in the normal, familiar course of travel without any scene change: course control (Wystrach et al. 2020 preprint). Adjustments, perhaps only small ones, to the left and right keep the traveler on the best course to head to its goal. The basic operation of the view-based navigational servomechanism in ants may be to command a turn to the left or turn to the right. Oscillating left and right keeps the ant on course in heading toward the ‘best’ view. If one keeps to a ruler-straight course of travel, one soon becomes uncertain as to whether the travel direction is still the best direction, there being a lack of comparisons with views in other directions (Cheng 2023). Managing uncertainty might be another function of oscillations.

### Conclusion

This study has demonstrated that desert ants oscillate as they navigate outbound and homebound. Their paths wiggle and their heads swing left and right. Both these oscillations, along with other behaviors, chief among them stopping to scan, are modulated in the face of scene changes. Scanning increases, the path meanders more, and with a large enough scene change, the amplitude of head swings and the spatial frequency of path oscillations increase. This entire suite of behaviors serves to increase visual exploration and thereby the learning of new visual cues.

## ACKNOWLEDGEMENTS

We acknowledge the traditional custodians of the land upon which this research was conducted, the Arrernte people. Their culture and customs have nurtured and sustained this land since the Dreamtime and continue to do so today. We pay our respects to their Elders past and present. We thank the Centre for Appropriate Technology at Alice Springs, Australia for letting us work on their property and providing some storage space, and the CSIRO Arid Zone Research at Alice Springs for administrative support. We are also thankful to Donald James McLean and Drew Allen for helping us with data analysis.

## Author contributions

Experimental design: SD. Data collection: SD; Data analysis: SD. Writing: SD and KC.

## Ethics standards

Australia has no ethical regulations regarding work with insects. The study was non-invasive and no long-term aversive effects were found on the nests or on the individuals studied.

## Data availability

Supplementary videos, Excel file and R scripts are available at Open Science framework: https://osf.io/6wk7a/files/osfstorage

---

**Table 1a.** Pearson correlations for Amplitude means and standard deviations (SD) in orientation oscillations in different conditions, along with their p-values.

| Amplitude mean and SD | Pearson correlation (r) | P Value |
| --- | --- | --- |
| C1 | 0.946 | < 0.0001 |
| +B | 0.924 | < 0.0001 |
| C2 | 0.942 | < 0.0001 |
| -B | 0.87 | < 0.0001 |

**Table 1b.** Pearson correlations for Temporal Frequency means and standard deviations (SD) in orientation oscillations in different conditions, along with their p-values.

| Temporal Frequency mean and SD | Pearson correlation (r) | P Value |
| --- | --- | --- |
| C1 | 0.33 | 0.24 |
| +B | 0.68 | 0.007 |
| C2 | 0.64 | 0.009 |
| -B | 0.81 | < 0.0001 |

**Table 1c.** Pearson correlations for Amplitude mean and standard deviations (SD) in path oscillations in different conditions, along with their p-values.

| Amplitude mean and SD | Pearson correlation (r) | P Value |
| --- | --- | --- |
| C1 | 0.94 | < 0.0001 |
| +B | 0.67 | 0.006 |
| C2 | 0.87 | < 0.0001 |
| -B | 0.88 | < 0.0001 |

**Table 1d.** Pearson correlations for Spatial Frequency means and standard deviations (SD) in path oscillations in different conditions, along with their p-values.

| Spatial Frequency mean and SD | Pearson correlation (r) | P Value |
| --- | --- | --- |
| C1 | 0.78 | 0.0005 |
| +B | 0.94 | < 0.0001 |
| C2 | 0.94 | < 0.0001 |
| -B | 0.82 | < 0.0001 |

**Table 1e.** Pearson correlations for Temporal Frequency means and standard deviations (SD) in path oscillations in different conditions, along with their p-values.

| Temporal Frequency mean and SD | Pearson correlation (r) | P Value |
| --- | --- | --- |
| C1 | 0.45 | 0.08 |
| +B | 0.64 | 0.008 |
| C2 | 0.5 | 0.05 |
| -B | 0.93 | < 0.0001 |

